# Using interactive platforms to encode, manage and explore immune-related adverse outcome pathways

**DOI:** 10.1101/2023.03.21.533620

**Authors:** Alexander Mazein, Muhammad Shoaib, Miriam Alb, Christina Sakellariou, Charline Sommer, Katherina Sewald, Kristin Reiche, Patricia Gogesch, Luise A Roser, Samira Ortega Iannazzo, Sapna Sheth, Susanne Schiffmann, Zoe Waibler, Vanessa Neuhaus, Susann Dehmel, Venkata Satagopam, Reinhard Schneider, imSAVAR Consortium, Marek Ostaszewski, Wei Gu

## Abstract

We address the need for modelling and predicting adverse outcomes in immunotoxicology to improve non-clinical assessments of immunomodulatory therapy safety and efficacy. The integrated approach includes, first, the adverse outcome pathway concept established in the toxicology field, and, second, the systems medicine disease map approach for describing molecular mechanisms involved in a particular pathology. The proposed systems immunotoxicology workflow is demonstrated with CAR T cell treatment as a use case. To this end, the linear adverse outcome pathway (AOP) is expanded into a molecular interaction model in standard systems biology formats. Then it is shown how knowledge related to immunotoxic events can be integrated, encoded, managed and explored to benefit the research community. The map is accessible online via the MINERVA Platform for browsing, commenting and data visualisation (https://minerva.pages.uni.lu). Our work transforms a graphical illustration of an AOP into a digitally structured and standardised form, featuring precise and controlled vocabulary and supporting reproducible computational analyses. Because of annotations to source literature and databases, the map can be further expanded to match the evolving knowledge and research questions.

## Introduction

Developing efficient tools for assessing the risks of immunomodulatory therapeutic modalities is a key step in improving predictivity of drug development during the non-clinical stage and offering innovative immunobiology models and biomarkers. The project imSAVAR (Immune Safety Avatar: nonclinical mimicking of the immune system effects of immunomodulatory therapies) aims at a better understanding of immunotoxic mechanisms and improving models (https://imsavar.eu). The adverse outcome pathway (AOP) concept is one of such tools that allows knowledge-based evaluation of the involved molecular mechanisms^1^.

In immunotoxicology, immune-related adverse outcome pathways (irAOPs) are used to visualise and study adverse effects of treatments. They allow highlighting a molecular initiating event (MIE), the key events (KEs), key event relationships (KERs) and an adverse outcome (AO), representing their order and also aligning these KEs to test systems and values of measured clinical parameters. These irAOPs can be used to help clinicians to assess the safety of a given treatment and propose new biomarkers and new treatment strategies^1–3^. Modelling strategies of AOPs are described in the OECD “Users’ Handbook supplement to the Guidance Document for developing and assessing Adverse Outcome Pathways”^4^. Specifically, the AOP graphical representation is discussed in the handbook in Development Tips 1 and 2^4^. A classical AOP is initially structured in a several-step linear diagram and stored as an image. While these graphical representations are informative and useful, there are limitations that make it difficult to interactively explore, share and use them for further modelling and prediction. We propose to address these limitations by employing advanced tools in systems biomedicine and network biology, e.g. the MINERVA platform^5^.

In systems biomedicine, the standardised representation and machine readability needed for interactive exploration, annotation and modelling of pathways are offered by approaches such as the disease maps. A disease map is a conceptual model of relevant mechanisms represented as a collection of interconnected signalling and metabolic pathways^6,7^. Examples of such maps are resources for cancer^8^, Parkinson’s disease^9^, rheumatoid arthritis^10^, asthma^11^ and, the most recent development, the COVID-19 Disease Map for capturing virus-host interaction mechanisms of the SARS-CoV-2 infection^12^. These maps can include multiple layers from molecular to intercellular and intertissue/interorgan interactions to reflect the physiological level of complexity^11^. Disease maps are designed for integrating prior knowledge, making sense of newly-generated data, modelling and predictions. The primary purpose of building a disease map is to structure knowledge about disease mechanisms in a single repository. The repository is usually integrated with a web-server-based user interface to enable interactive visualisation and exploration of this knowledge. State-of-the-art disease maps also utilise the systems biology standards to ensure interoperability with other biological resources.

In both approaches, irAOPs and disease maps, the basis is a pathway representing key events that result in a particular outcome. The approaches differ and complement each other in the level of details and in their main focus. The irAOP focuses on the higher level of molecular and physiological events, multiple levels of entities (key molecules, cells, tissues, organs, organism), a variety of functional relationships (switch on, switch off, branching), as well as quantitative clinical parameters and assays. They have a common structure consisting of a MIE, a series of KE connected by KERs and an AO. They usually do not provide a comprehensive molecular description of every aspect of a biological process *per se*. The disease map, on the other hand, concentrates on detailed molecular mechanisms relevant to induce a particular condition and has the ability to visualise the higher-level relationships required in immunotoxicology. Integrating the concepts of immunotoxicology irAOP with the systems biomedicine disease map offers a promising improvement to the classical irAOP.

By combining the two approaches, we aim to build a solution that combines their advantages while addressing anticipated knowledge gaps. It also paves the way to tackle the challenges of multi-scale representation including molecular, cellular and immune system levels, with a perspective of creating an executable computational model for making predictions.

Major advantages of the enhanced pathway-based AOP approach compared to the standard modelling and visualisation frameworks as recommended by the OECD AOP concept^4^ and the AOP Wiki (https://aopwiki.org/aops): 1) focus on relevant molecular pathways with an ability to represent intercellular and physiological relationships; 2) applying well-established standards and editors for the reconstruction of the underlying biology; 3) identifying knowledge gaps during the reconstruction process and clarifying the mechanisms of the MIE, KEs, KERs and AO; 4) using the MINERVA platform for online visualisation and exploration, with such entities as proteins, genes and metabolites identified and linked to appropriate external databases, and with multiple plugins available. In more details the approach and the MINERVA platform are discussed in the Methods section.

In this work, we propose an integrated systems toxicology framework and create a proof-of-concept knowledge-based and irAOP-based map of molecular interactions for the cytokine release syndrome mediated by CAR T cells.

## Methods

### The representation of the underlying biology for adverse outcome pathways

The irAOP concept connects the MIE and the corresponding AO via a series of KEs and KERs known to be involved at different levels of biological organisation^1,4^. In one of the following works, a linear adverse outcome pathway was expanded into a complex network of molecular events, with feedback and feedforward loops and inter-relationships between individual key events presented^2,3^. By building on this example, we applied the state-of-the-art advances in the standard systems biology field and used such formats as the Systems Biology Graphical Notation (SBGN)^13^ and Systems Biology Markup Language (SBML)^14^ to formalise and visualise our knowledge.

### The systems biology approach for adverse outcome pathways

In systems medicine, a disease map is a conceptual model of disease mechanisms, with events depicted on the level of molecular interactions^6,7^. Figure 1 describes the systems immunotoxicology framework integrated from the adverse outcome pathways concept and the adapted systems medicine disease map methodology^15^.

**Figure 1.**
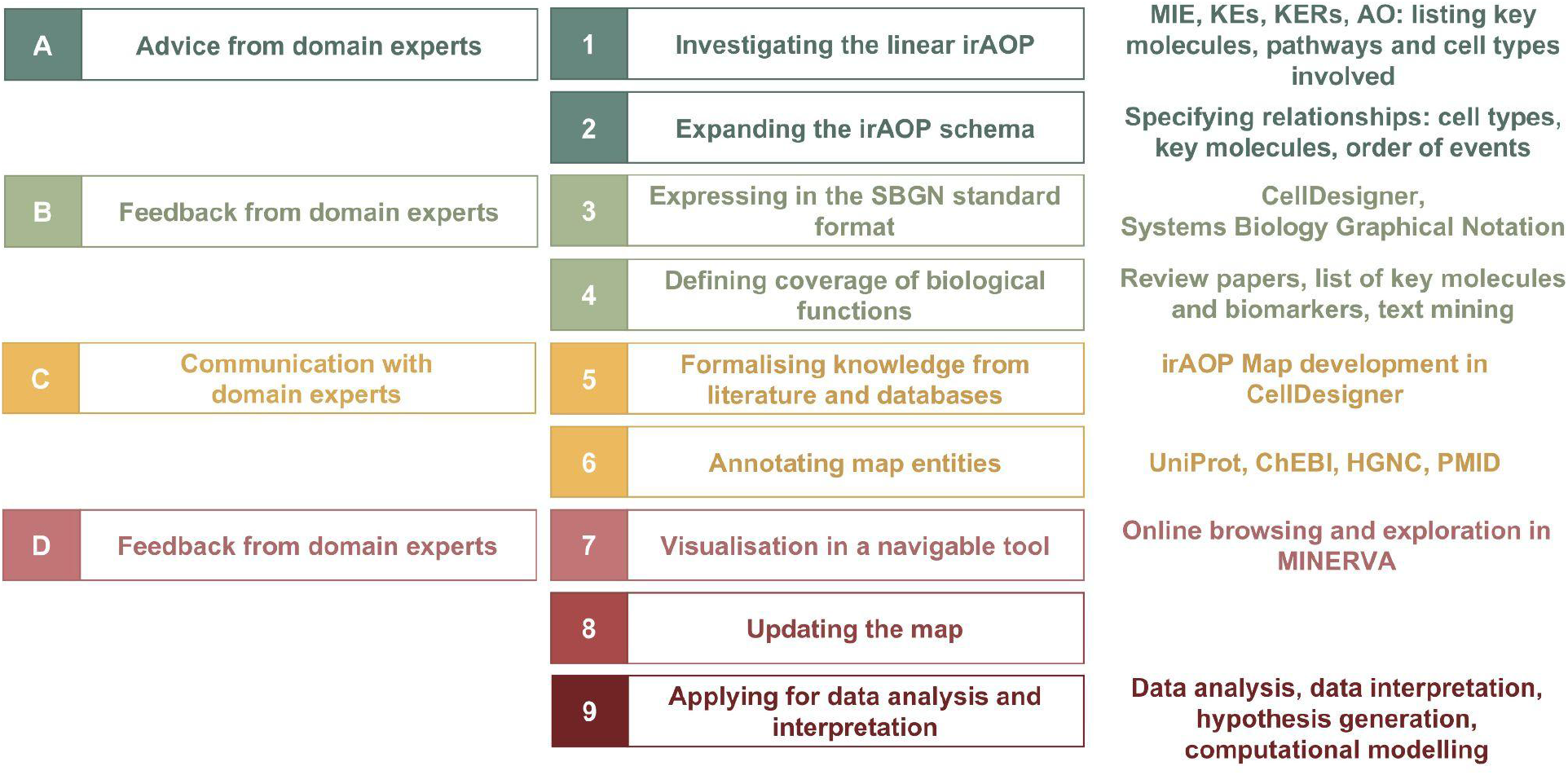
The systems framework for knowledge management and exploration in immunotoxicology. irAOP, immune-related adverse outcome pathway; MIE, molecular initiating event; KEs, key events; KERs, key event relationships; AO, adverse outcome; UniProt, a database of protein sequence and functional information (https://www.uniprot.org); ChEBI, Chemical Entities of Biological Interest (https://www.ebi.ac.uk/chebi); HGCN, HUGO Gene Nomenclature Committee names (https://www.genenames.org); PMID, references in the form of PubMed IDs (https://pubmed.ncbi.nlm.nih.gov). Descriptions of the steps are provided in the text.

Steps to construct the irAOP maps:

*Step 1. Investigating the linear irAOP*. Key molecules, related pathways and cell types involved are hypothesised. The relevant publications are reviewed. The information is organised in the form of a weight-of-evidence-table for all the key events and discussed with the experts (Alb et al., in preparation).
*Step 2. Expanding the irAOP schema*. A top-level view is built by representing key biological mechanisms and connecting the previously identified key molecules into a network.
*Step 3. Expressing knowledge in the standard SBGN format*. When the network components are identified and connected, the biological mechanisms are expressed in the standard graphical systems biology languages^13^ to have both a human- and computer-readable top-level view diagram, thus ensuring shareability and enabling computational modelling.
*Step 4. Defining coverage of biological functions*. The boundaries of the conceptual model are assessed and relevant key pathway modules are listed as the main components of the map.
*Step 5. Formalising knowledge from literature and databases*. Based on the top-level view (Steps 2 and 3) and our assessment of the complexity (Step 4), a pathway-level detailed network of molecular interactions is built. Systems biology editors are employed for the network construction, e.g. CellDesigner (http://www.celldesigner.org), Newt Edito^r16^ (http://newteditor.org), SBGN-ED^17^ (http://sbgn-ed.org) or the yEd Graph editor (https://www.yworks.com/products/yed) in combination with the ySBGN converter (https://github.com/sbgn/ySBGN).
*Step 6. Annotating the map entities*. The map objects have to be identified by linking them to external databases. For example, proteins are identified via UniProt IDs (https://www.uniprot.org) and molecular interactions are confirmed with PubMed IDs (https://pubmed.ncbi.nlm.nih.gov). For that, we follow stable identifiers using the Identifiers.org^18^ or MIRIAM annotation^19^.
*Step 7. Visualisation in a navigable tool*. For making the results easily accessible and explorable we use the MINERVA platform^5,20,21^ (see the next Section for details).
*Step 8. Updating the map*. With newly published information or feedback from the research community, the map can be further refined and updated. Modularised structure makes it easier to manage and update the map components^22^.
*Step 9. Applying for data analysis and interpretation*. The complete map can be used for ‘omics data visualisation, analysis and interpretation^12^. The map can also be transformed into a simulatable computational model, for example, a Boolean model, for predictions and advanced hypothesis generation^10^.

Communication with domain experts (A, B, C and D in Figure 1) is one of the most important components ensuring adequate representation of knowledge from published research on the topic, prioritising key aspects and making the map a practically applicable resource.

### The MINERVA Platform

The MINERVA Platform is a standalone web-service for user-friendly online visualisation and exploration of extensive systems biology networks^5^. The MINERVA documentation is available at https://minerva.pages.uni.lu.

The MINERVA Platform allows interactive browsing of biological networks, provides access to their annotation and connects to external databases for more information about the molecules involved. Evidence for interactions is stored as PubMed IDs with links to PubMed entries. Custom data can be uploaded and visualised in colours of different intensities according to the values provided. MINERVA enables search and exploration of drug targets via online queries to DrugBank^23^ (https://www.drugbank.com) and ChEMBL^24^ (https://www.ebi.ac.uk/chembl).

MINERVA works with CellDesigner XML format, SBGN and SBML with layout information.

Video demo tutorials are available for asthma as a use case^11^ and cover such functionalities as 1) navigating and searching in MINERVA, 2) adding comments directly on the map, 3) exploring the drug target search functionality, and 4) visualising ‘omics data (https://asthma-map.org/tutorials).

## Results

### Immune-related adverse outcome pathway of CAR T cell treatment

Cytokine release syndrome (CRS) is the most common type of toxicity caused by CAR T cell therapy and, in general, is one of the common types of toxicity caused by advanced therapy medicinal products (ATMPs)^25–28^. Figure 2a presents the proposed respective irAOP. The MIE, the KE and the AO are described below.

**Figure 2.**
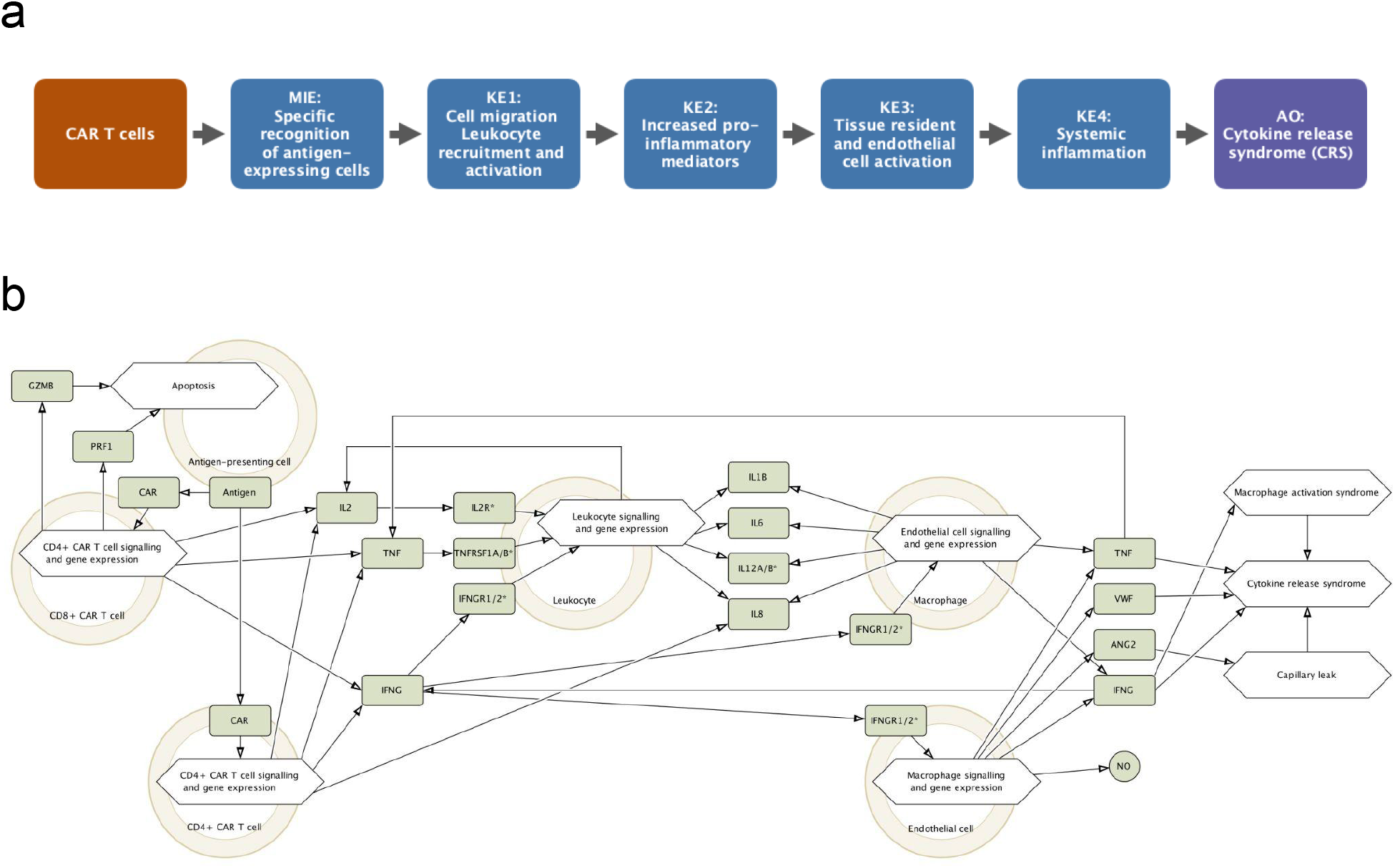
CAR-T-cell-induced cytokine release syndrome (CRS): a comparison of a linear irAOP and a more detailed representation of the underlying biology. **a.** The linear irAOP for the CAR-T-cell-induced cytokine release syndrome. MIE, molecular initiating event; KE, key event; AO, adverse outcome. **b.** The top-level view of the CAR T cell irAOP-Map - the underlying biology. The map is available in MINERVA for commenting and exploring at https://imsavar.elixir-luxembourg.org/minerva/index.xhtml?id=cart14.

#### Molecular initiating event (MIE): specific recognition of antigen-expressing cells

Expression of a chimeric antigen receptor (CAR) enables T cells to recognise and bind to antigen-expressing cells such as tumour cells in a non-MHC restricted manner. As a result, CAR T cells are activated^29–32^.

#### Key event 1 (KE1): activation of CAR T cells

The activation of CD8^+^ CAR T cells leads to the release of cytolytic enzymes such as granzyme B (GZMB), and both activated CD8^+^ and CD4^+^ CAR T cells produce cytokines including IL-2 and IFN-γ^32–35^.

#### Key event 2 (KE2): increased proinflammatory mediators

The released cytokines can be measured by using cytokine release assays and multiplex analysis tools. The production of IFNG by CAR T cells triggers IL-6 production by monocytes and endothelial cells and IL-1, IL-6 and nitric oxide by macrophages^38–40^.

#### Key event 3 (KE3): tissue-resident and endothelial cell activation

The increased levels of proinflammatory mediators then lead to inflammation and its amplification, as well as to activation of other immune cells. This means enhanced IL-6 and IL-1 production by monocytes and activation of endothelial cells with production of von Willebrand factor (VWF) and angiopoietin-2 (ANG2)^28^. VWF plays a key role in coagulation^41^, and ANG2 promotes capillary leak^42^.

#### Key event 4 (KE4): systemic inflammation

Consequently, systemic inflammation develops. It includes an increase of pro-inflammatory cytokines and activation of the innate immune system. The levels of IL-6 and C-reactive protein are inflammatory markers of systemic inflammation.

#### Adverse outcome (AO): cytokine release syndrome

CRS is linked to macrophage activation syndrome which is associated with activated lymphocytes that produce IFNG and activated macrophages that produce IL-6 and TNF-α^37,39,40,43^.

This knowledge was formalised into a top-level overview diagram with the key molecules mapped and interconnected (see below).

### The extended representation of the underlying biology

Based on the linear AOP and the relevant textual information, we built the top-level view of the CAR T cell irAOP-Map. For that, we analysed the available information. While in the textual description to the linear irAOP the information was sufficient, it needed to be extended for the requirements of the biological network construction. When reviewing known molecules involved, we identified cases of non-specific proteins: IL-1 means two different proteins IL1A and IL1B, and NFAT can potentially mean five different proteins in UniProt (Table 1, columns “Symbol” and “Identifier”). The analysis of the source and target cell types helped to find connectivity in the network and identify the gaps in the current model (Table 1, columns “Source” and “Target”). The first version of the top-level view of the CAR T cell irAOP-Map is shown in Figure 2b.

**Table 1.** CAR T treatment: key molecules involved, with HGNC symbols and UniProt identifiers. n/a, not applicable.

| Name | Symbol | Identifier | Source | Target | Receptors (candidates) |
| --- | --- | --- | --- | --- | --- |
| IL-2 | IL2 | UniProt:P60568 | CAR T cell, Leukocyte | Leukocyte | IL2RA, IL2RB, IL2RG |
| IFN-γ | IFNG | UniProt:P01579 | CAR T cell, Macrophage, Endothelial cell | Leukocyte, Macrophage, Endothelial cell | IFNGR1, IFNGR2 |
| TNF-α | TNF | UniProt:P01375 | CAR T cell, Macrophage, Endothelial cell | Leukocyte | TNFRSF1A, TNFRSF1B |
| CD69 | CD69 | UniProt:Q07108 | CAR T cell | Antigen-expressing cell (via ligand) | n/a |
| CD25 | IL2RA | UniProt:P01589 | CAR T cell | n/a | n/a |
| NFAT | NFATC1, NFATC2, NFATC4, NFATC3, NFAT5 | omitted | n/a (transcription factors) | n/a (transcription factors) | n/a (transcription factors) |
| Perforin | PRF1 | UniProt:P14222 | CAR T cell | Antigen-expressing cell | n/a |
| Granzyme B | GZMB | UniProt:P10144 | CAR T cell | Antigen-expressing cell | n/a |
| IL-1 | IL1A, IL1B | UniProt:P01583; UniProt:P01584 | Leukocyte, Macrophage | T cell, Neutrophil, B cell | IL1R1, IL1R2 |
| IL-6 | IL6 | UniProt:P05231 | Leukocyte, Macrophage | T cell | IL6R, IL6ST |
| NO | nitric oxide | CHEBI:16480 | Macrophage | Tumor cell | n/a |
| IL-8 | CXCL8 | UniProt:P10145 | CAR T cell | T cell | CXCR1, CXCR2 |
| IL-12 | IL12A; IL12B | UniProt:P29459; UniProt:P29460 | CAR T cell, Leukocyte, Macrophage | T cell | IL12RB1, IL12RB2 |
| C-reactive protein | CRP | UniProt:P02741 | Hepatocytes, macrophages, endothelial cells, lymphocytes | n/a (clinical parameter) | n/a (clinical parameter) |
| Ferritin | FTH1; FTL; FTMT | UniProt:P02794; UniProt:P02792; UniProt:Q8N4E7 | Macrophage | n/a (clinical parameter) | n/a (clinical parameter) |
| MCP1 | CCL2 | UniProt:P13500 | Macrophage | Endothelial cell | CCR2 |
| IL-8 | CXCL8 | UniProt:P10145 | Leukocyte | Neutrophil | CXCR2 |
| VWF | VWF | UniProt:P04275 | Endothelial cell | Platelets | P1BA, GP1BB |
| ANG2 | VPS51 | UniProt:Q9UID3 | Endothelial cell | Endothelial cell | TEK |

Table 1 demonstrates how network reconstruction helps to ask questions and identify gaps in our understanding of the biological processes involved. We clarify and formalise available knowledge by identifying molecules and specifying the listed cell types. In such a map development protocol, by necessity, for ensuring the connectivity between the map objects, we have to actively investigate the connections, the receptors and cell types involved. For example, knowing that IL-2 is produced by CAR T cells and affects leukocytes, as the next step, we focus on IL-2 receptor proteins IL2RA, IL2RB and IL2RG, the corresponding signalling cascade and the activated transcription factors (Figure 3).

**Figure 3.**
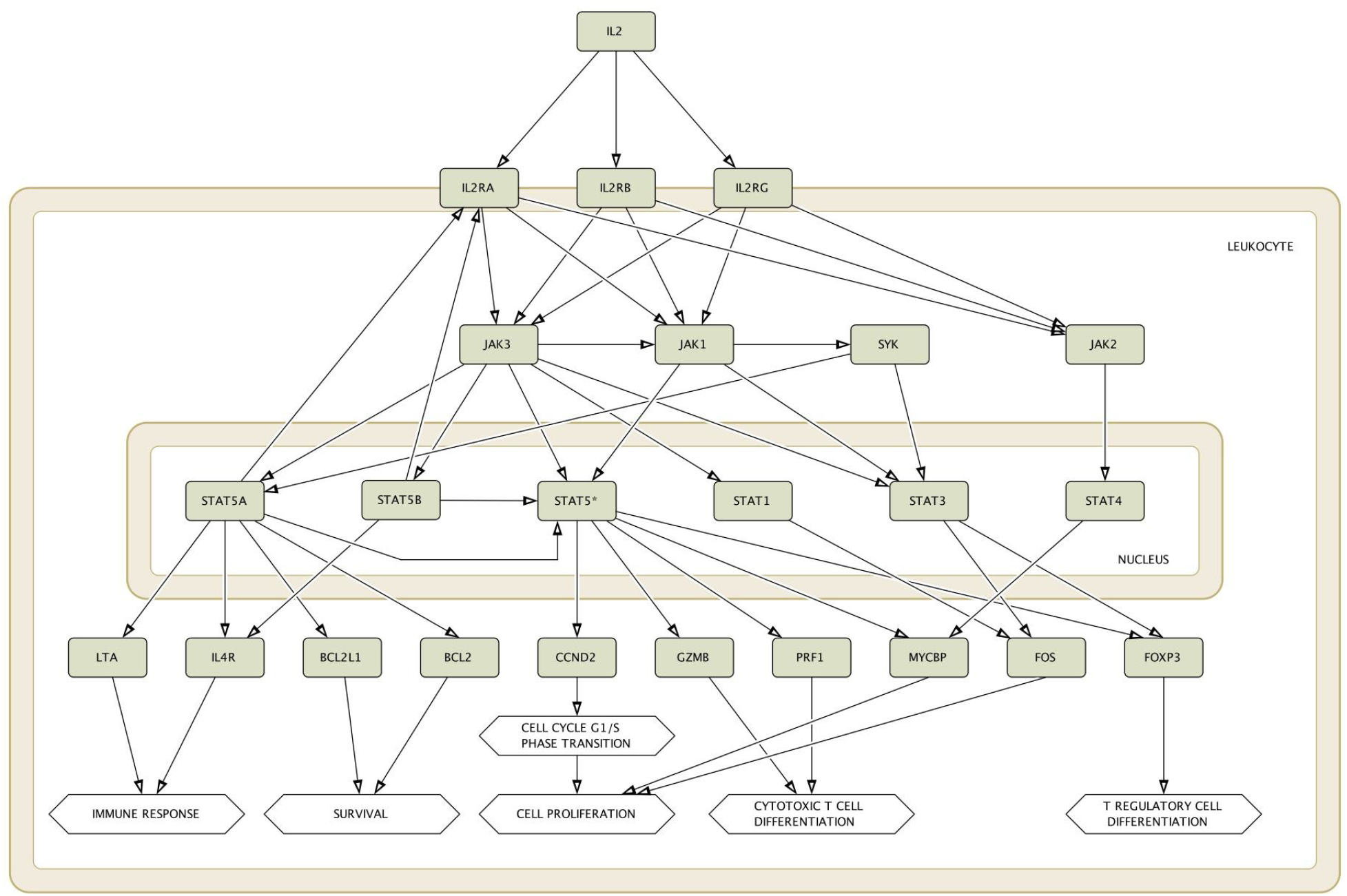
IL-2 signalling. This pathway is manually converted to the SBGN Activity Flow format from the MetaCore pathway map ID 2770 Immune response IL-2 signalling via JAK-STAT (https://portal.genego.com). The proteins are presented with the use of HGNC names (https://www.genenames.org). The map is available in MINERVA for commenting and exploring at https://pathwaylab.elixir-luxembourg.org/minerva/index.xhtml?id=il2v09.

### Future development

Within the imSAVAR project (https://imsavar.eu), we plan to extend the CAR T cell irAOP-Map, refine the described systems immunotoxicology protocol and demonstrate the applicability of such knowledge databases resources. One of the promising applications is developing executable computational models for predictions and hypothesis generation^44,45^, potentially with the use of the CaSQ tool^46^ and the Cell Collective Boolean modelling pipeline^47–49^.

## Conclusion

In the proposed framework, we bring together the recent advances in the immunotoxicology and systems biomedicine fields. We bridge the experience of developing conceptual descriptions of disease mechanisms, disease maps, to the practice of describing immunotoxic effects in the form of AOPs. With this, the AOP concept is enhanced with detailed description of intracellular and intercellular molecular interactions captured in the standard systems biology format. Focusing on the CAR-T-cell-treatment-induced CRS, we demonstrate how a molecular interaction map can be developed from a linear AOP and offer the results in an easily accessible and explorable way in the web-based MINERVA platform. On the other hand, this framework can improve the development of disease-specific AOPs, identify knowledge gaps in existing AOPs and lead to the design of new experiments. Because we apply a transparent and consistent methodology with the use of standards, this framework can be reused and applied to other immunomodulatory treatment scenarios. Continuing our work in this direction we aim to build explorable knowledge resources that could benefit immunotoxicology research and contribute to improving non-clinical assessment of immunomodulatory therapies.

## Key points

- In immunotoxicology, an adverse outcome pathway shows a sequence of molecular and cellular events that result in a toxic outcome upon treatment with a specific drug.
- In systems biomedicine, a disease map is a description of disease mechanisms on the levels of molecular interactions and intercellular communication for integrating prior knowledge, making sense of newly-generated data, modelling and predictions.
- We are applying the disease map approach to the area of immunotoxicology and offer an interactive web-based platform for expanding immune-related adverse outcome pathways to detailed representations of the underlying biology.
- The objective is to model adverse outcomes as a non-clinical assessment strategy by integrating our understanding of the disease complexity and knowledge on the mechanisms of the adverse outcomes of the treatment.
- We focus on the adverse outcome pathway of CAR T cell treatment and from a simplified linear pathway build a detailed representation of the underlying biology.

## Abbreviations

AO: adverse outcome
ATMPs: advanced therapy medicinal products
CAR: chimeric antigen receptor
CRS: cytokine release syndrome
irAOP: immune-related adverse outcome pathway
KE: key event
KERs: key event relationships
MIE: molecular initiating event
SBGN: Systems Biology Graphical Notation
SBML: Systems Biology Markup Language

## Acknowledgements

We would like to acknowledge the Responsible and Reproducible Research (R3) team of the Luxembourg Centre for Systems Biomedicine for supporting the project. The work presented in this paper was carried out using the ELIXIR Luxembourg tools and services.

## Data availability statement

All the maps are freely accessible online via the MINERVA platform.

